# Structure-Guided Discovery of CHI3L1 Inhibitors from Ultralarge Chemical Spaces for Glioblastoma Therapy

**DOI:** 10.1101/2025.08.09.669500

**Authors:** Baljit Kaur, François Sindt, Longfei Zhang, Didier Rognan, Moustafa Gabr

## Abstract

Glioblastoma (GBM) is the most aggressive primary brain tumor, with a median survival of approximately one year and limited therapeutic options. Chitinase 3-like 1 (CHI3L1) is increasingly recognized as a promising target in GBM due to its role in tumor progression and immune modulation. In this study, we employed an in-house structure-based screening strategy (SpaceDock) to explore a virtual chemical space of 377 billion compounds for potential CHI3L1 inhibitors. Using a reaction-aware ligand design approach, 60 top-scoring virtual hits were synthesized, and 45 were obtained with sufficient purity for experimental testing. Primary screening by microscale thermophoresis (MST) identified nine hits. Compounds showing dose-dependent binding were subsequently analyzed using surface plasmon resonance (SPR), leading to the identification of compound **9e** with a dissociation constant (*K*_d_) of 19.11 µM. Importantly, **9e** demonstrated robust, dose-dependent efficacy in a multicellular 3D GBM spheroid model, significantly reducing spheroid viability and downstream STAT3 signaling. These results highlight **9e** as a promising drug candidate for modulating the CHI3L1–STAT3 axis and underscore the potential of structure-guided, reactivity-aware virtual screening of ultralarge chemical spaces to target non-enzymatic, conformationally dynamic proteins in complex cancer models.

---

Glioblastoma (GBM) is an aggressive and notoriously hard-to-treat brain cancer.^1–7^ Its ability to spread extensively and resist conventional therapies is made worse by the presence of a highly variable cell population and continuous interaction with its surrounding environment.^8–12^ Although commonly used treatments such as surgery, radiation, and chemotherapy form the backbone of clinical care,^13–15^ they often fail to significantly improve patient outcomes. Each year, this disease is responsible for over 10,000 deaths in the United States alone,^16^ and most patients survive only about one to one and a half years with only a quarter surviving beyond the first year and very few reaching five years after diagnosis.^17–19^

What sets GBM apart from many other cancers is its ability to reshape the immune landscape in its favor.^20–22^ Rather than prompting a defensive reaction, the tumor reprograms immune components to promote its own growth and survival. A key part of this process involves manipulating immune cells especially macrophages into supporting the tumor instead of fighting it. These altered immune cells, driven by complex signaling pathways, help the tumor grow, build blood vessels, and suppress further immune activity.^23–25^ This reprogramming of the immune microenvironment not only speeds up disease progression but also makes current immune-based treatments less effective. As a result, there is a pressing need to discover new molecular targets that could help reverse this immune suppression.^26,27^

Even with the use of approved drugs like temozolomide,^28,29^ bevacizumab,^30^ and emerging approaches such as tumor-treating fields,^31–33^ mutations in the IDH1^34,35^ or EGFR genes^36,37^ efforts to treat GBM remain largely unsuccessful. Major barriers include the tumor’s resistance to treatment, the challenge of delivering drugs across the blood-brain barrier, and the immunosuppressive environment within the tumor itself.^38–42^ These persistent difficulties highlight the urgent need for new small-molecule drugs and better screening methods that can lead to more effective therapies.

Among the potential molecular targets receiving growing attention is Chitinase-3-like protein 1 (CHI3L1), also known as YKL-40.^43–46^ This protein is secreted by cells^47–51^ and, although it shares structural similarities with active enzymes called chitinases, it lacks the key residues needed for enzymatic function.^52,53^ Instead, it plays a significant role in processes like inflammation,^54–57^ tissue remodeling,^58–60^ and cancer progression.^61–63^ In the case of GBM, CHI3L1 levels are often elevated, and the protein appears to help steer macrophages toward a state that supports tumor growth, promotes new blood vessel formation, and suppresses immune responses-all of which create a tumor-friendly environment.^44,64,65^ Through these connections, CHI3L1 affects numerous signaling cascades involved in tumor biology and immune evasion. Notably, it has been implicated in the activation of the STAT3 pathway, a key driver of immunosuppression and tumor cell survival in GBM. ^64,65^

Despite this compelling biological rationale and clinical association, the pursuit of selective small-molecule inhibitors targeting CHI3L1 has remained limited.^66–68^ What makes CHI3L1 both intriguing and difficult to target is that it doesn’t operate like a traditional drug target. It doesn’t rely on enzymatic activity or single receptor-ligand interactions. Rather, it functions as a flexible and context-specific signaling regulator, interacting transiently with several surface receptors such as IL-13Rα2 (IL-13 receptor alpha 2), RAGE (receptor for advanced glycation end products), and VEGFR2 (Vascular Endothelial Growth Factor Receptor 2).^69,70^ Through these connections, CHI3L1 affects numerous signaling cascades involved in tumor biology and immune evasion.^71,72^ Because of its multifunctional nature and lack of catalytic activity, CHI3L1 exemplifies a class of biologically crucial yet pharmacologically elusive targets. Tackling such proteins will require innovative therapeutic strategies that move beyond conventional drug design, offering new opportunities for more effective GBM treatments.

We herewith present one of the largest virtual screening approaches ever described, leveraging the efficiency of our recently described SpaceDock virtual screening method,^73^ to search in a chemical space of 377 billion synthetically feasible molecules for CHI3L1 inhibitors. Among the 45 candidates that could be synthesized, three compounds could be validated by several orthogonal assays and appear as promising hits for further optimization.

## Computational and experimental methods

### Docking of chemical reagents to human chitinase-3-like protein 1 (CHI3L1)

The X-ray structure of the human CHI3L1 in complex with the inhibitor 30^68^ was downloaded from the Protein Data Bank (PDB 8R4X). Hydrogen atoms and simultaneous optimization of protonation states of protein, water and ligand atoms was performed with Protoss v.4.0.^74^ All organic molecules (water, chloride ions, inhibitor) were removed keeping only remaining protein atoms of chain A which were saved in mol2 file format. The commercial building blocks were chosen from a pool of 183,981 reagents utilized in the REAL Space by Enamine (2022q3-4 update). The cavity was detected from X-ray atomic coordinates of CHI3L1-bound inhibitor 30. Up to 20 docking poses were generated for each reactant using default settings of GOLD v.2024.1.0^75^ and scored by the PLP scoring function.

### Ligand enumeration by reagents coupling (SpaceDock)

Given pairs or triplets of chemically compatible reagents poses, a ligand was generated within the protein binding site, according to their respective location and chemical compatibility, as previously described.^73^ Reagent poses were initially loaded using an in-house mol2 parser and annotated for 39 two-component and one three-component organic chemistry reactions (**Table S1**). This process was repeated for each reaction, following a similar workflow. A subsequent set of filters was then applied to pairs of reagent poses, including the distance between their center of mass to promptly eliminate distant pairs, the distance between connectable atoms, examination of certain angles of the future formed bond/ring to ensure a suitable geometry, and consideration of clashes (≤4 between non-reacting atoms) to prevent overlapping substituents. If a pair of reactants satisfied all the rules, a bond/ring closure was created between the connectable atoms. The hybridization of reacting atoms was then updated to reflect the newly created bonds and exit atoms (to be removed after the reaction) were deleted. For each of the 415,064,938 valid recombinations, the fully enumerated molecule was then saved into a single mol2 file. To reduce the number of solutions, only recombinations forming a hydrogen bond to key Y206 and D207 residues,^68^ detected with IChem v.5.2.8,^76^ were kept. Despite this filter, the number of solutions remained high (130,474,687). A drug-likeness filter was then applied to the enumerated ligands, as previously described^77^ to keep only 61,061,738 drug-like molecules. The remained recombinations, in presence of the target protein, were last energy-minimized in Szybki v2.4.0.0 (Openeye Scientific Software, Sante Fe, NM, U.S.A.) using standard settings and the MMFF94 force-field.^78^ Only minimized poses whose heavy atoms deviated by less than 1 Å root-mean-square (rms) deviations to the native non-minimized SpaceDock poses were kept to retain 41,367,191 plausible solutions. Finally, a last verification was performed using IChem v5.2.8 to ensure that key hydrogen bonds with Y206 and D207 were preserved after minimization. Additionally, two aromatic interactions with F261 and W352 were checked, resulting in a final set of 2,680,769 minimized SpaceDock poses.

### Redocking of SpaceDock poses

The coupling of two reagent poses, followed by protein constraint refinement (referred to as the “SpaceDock” pose), was redocked into the target protein structure using GOLD. The scoring function employed was PLP, with 20 generated poses, and the same parameter file as previously. To be retained for subsequent investigations, a ligand had to fulfil several criteria: (i) rmsd to the minimized SpaceDock pose less than 2.0 Å, (ii) hydrogen-bond to Y206 and D207, (iii) aromatic interactions to F261 and W352. If multiple docking poses from the same ligand satisfied these rules for each SpaceDock pose, all of them were retained, leaving 1,074,957 poses for further evaluations.

### Quality check of redocked poses

The number of torsion strains in every redocking pose was estimated with TorsionAnalyzer v.2.0.0.^79^ Any pose with at least one torsion annotated as ’strained’ was discarded from further analysis. Local strain (distortion of the specific conformation from the nearest local minima) and global strain (energy required to select the specific conformation from the full conformational ensemble of the corresponding compound in implicit water) energies were then computed with standard parameter of Freeform v.2.4.0.0 (Openeye Scientific Software, Sante Fe, NM, U.S.A) Any pose with local and global strain energies higher than 4 and 8 kcal/mol, respectively, were discarded. Last, remaining poses were inspected, in their protein-bound state, for counting the number of unsatisfied ionic bonds, hydrogen-bond donors and acceptors. First, protein-ligand ionic and hydrogen-bonds were registered with IChem. Any charged atom or hydrogen-bond donor/acceptor atom of the ligand (according to IChem definitions)^80^ not present in the above list was annotated as “unsatisfied” atom. Unsatisfied heavy atoms being both donors and acceptors (e.g. hydroxyl oxygen atom) were only counted once. Ligand atoms participating to intra-molecular hydrogen bonds were considered as satisfied. Altogether, ligand poses with more than 2 unsatisfied donors and 4 unsatisfied acceptors were removed from the final hit list. A total of 77,605 poses was saved for the next rescoring step.

### Commercial availability and rescoring

Commercial availability in REAL space (Enamine Ltd, Kiyv, Ukraine) was checked with the Enamine REAL space API (https://real.enamine.net). All available hits were rescored with two rigorous physics-based scoring functions: Hyde v.2.1.1^81^ and VM2^82^. Since multiple solutions may still be valid for the same ligand, ligand redundancy was removed by keeping the best Hyde score for each ligand. Any pose exhibiting a predicted Hyde affinity below 1µM and a VM2 predicted absolute binding free energy less than 6 kcal.mol^-1^ were saved. A total of 1,794 ligands fulfilled these two criteria

### Final hit selection

The 1,794 unique ligands were clustered by maximum common substructure (MCS) using ChemAxon LibMCS program (https://docs.chemaxon.com/display/docs/jklustor_library-mcs-libmcs-clustering.md) and a minimal MCS size of 9 atoms. For each of the 138 clusters, the best VM2 score was considered as representative. The best 60 ligands according to their VM2 predicted absolute binding free energy were selected to define the final hit list, whose synthesis was purchased at Enamine. Out of the 60 requests, 45 ligands (**Table S2**) could be synthesized as LC-MS pure compounds (UV-vis purity > 90%).

### Microscale Thermophoresis (MST) single dose screening

To evaluate protein–ligand interactions, CHI3L1-His protein was labeled using a RED-tris-NTA fluorescent dye (NanoTemper Technologies) following the kit protocol. The labeling reaction was performed by incubating 200 nM of the protein with 100 nM dye in PBS containing 0.05% Tween-20 for 30 minutes in the dark at room temperature. After labeling, the protein was diluted in assay buffer (10 mM HEPES, 150 mM NaCl, 0.1% Pluronic F-127, 1 mM TCEP, and 8% DMSO (pH 7.4)) and combined with each test compound to a final concentration of 20 nM protein and 250 μM compound. Samples were incubated at ambient temperature for 30 minutes, centrifuged briefly, and transferred to the Dianthus NT.23 Pico system for analysis. Controls included buffer with DMSO alone. All measurements were performed in triplicate, and average values were reported. To assess background fluorescence, test compounds at 250 μM were prepared from DMSO stocks in assay buffer, incubated in the dark, centrifuged, and measured for signal using the same platform. Fluorescence quenching was also evaluated by mixing 10 nM dye with each compound under similar conditions. All assays were repeated three times, and results expressed as mean ± standard deviation.

### Dose–Response MST Analysis

For compounds identified as primary hits, binding affinities were further examined using serial dilutions (16-point range from 500 uM to low nM) mixed with the labeled CHI3L1-His protein. The final DMSO concentration was adjusted to 4%. After 30 minutes of incubation, samples were loaded into MST capillaries, and measurements were taken using the Monolith NT.115 instrument. Settings included moderate to high IR-laser power and 60–80% LED intensity in the red channel. Data were analyzed using MO.Affinity Analysis software (NanoTemper).

### Surface Plasmon Resonance (SPR)

SPR experiments were conducted on the Biacore 8K platform to confirm binding between CHI3L1-His and confirmed hits. The protein was immobilized on a sensor chip and increasing concentrations of the compound were injected sequentially using a single-cycle kinetic format. All assays were carried out at 25 °C. Reference subtraction was performed using signals from a blank flow cell and buffer-only injections. Binding kinetics were analyzed using Biacore™ Insight Evaluation Software.

### AlphaLISA-Based Inhibition Assay Protocol

Tested compounds were prepared in assay buffer and incubated with hCHI3LA-His (Cat# CH1-H5228, Acro Biosystems, Newark, DE, USA) and Galectin-GST (Cat# 10289-H09E, Sino Biological, Beijing, China) at final concentrations of 100 μM (compound), 60 nM (hCHI3LA-His), and 60 nM (Galectin-GST). The mixture was incubated at room temperature (r.t.) for 3 hours. Subsequently, AlphaLISA anti-6His donor beads (AS116D, Revvity, Waltham, MA, USA) and anti-GST acceptor beads (AL110C, Revvity) were added, followed by an additional 1-hour incubation at r.t. in the dark. Signal detection was performed using a Tecan Infinite M1000 Pro plate reader set to AlphaLISA mode. The assay was conducted in technical triplicate, and results are reported as mean ± standard deviation (SD).

### Cell viability of glioblastoma (GBM) spheroids

The GBM spheroids were prepared as previously described.^83^ U-87 MG GBM cells (ATCC, Cat# HTB-14) were cultured in DMEM (ATCC, Cat#30-2002) containing 4.5 g/L glucose and 2 mM L-glutamine, supplemented with streptomycin and penicillin. HMEC-1 (ATCC) were maintained in MCDB131 medium with 10 mM L-glutamine, 10 ng/ml FGF, and 1 µg/ml hydrocortisone. Briefly, U-87 MG and HMEC-1 cells were co-seeded with macrophages on low-adhesion 96-well plates at 2 × 10³ cells per well in 100 µl of the tested compound at varying concentrations (25, 50, and 100 µM) or control media. After 72 hours, cell viability was assessed using the CCK-8 assay (MedChemExpress, Cat# HY-K0301) according to the manufacturer’s recommended protocol. Absorbance at 450 nm was measured using a microplate reader.

### Homogeneous time-resolved fluorescence (HTRF) assay

Assessment of phospho-STAT3 levels in GBM spheroids was determined by HTRF phospho-STAT3 kit from Revvity (Cat# 62AT3PET) using the manufacturer’s recommended protocol. All experiments were conducted in triplicate.

## Results and Discussion

### Structure-based screening of a virtual chemical space of 377 billion compounds

We here applied an in-house structure-driven, reaction-aware ligand design strategy (SpaceDock)^73^ tailored to the unique demands of non-enzymatic, structurally flexible proteins such as CHI3L1. Moving beyond reliance on extensive pre-assembled virtual libraries, our approach initiates from commercially available building blocks, which are docked onto experimentally or computationally characterized protein surfaces of interest. (**Figure 1**).

**Figure 1.**
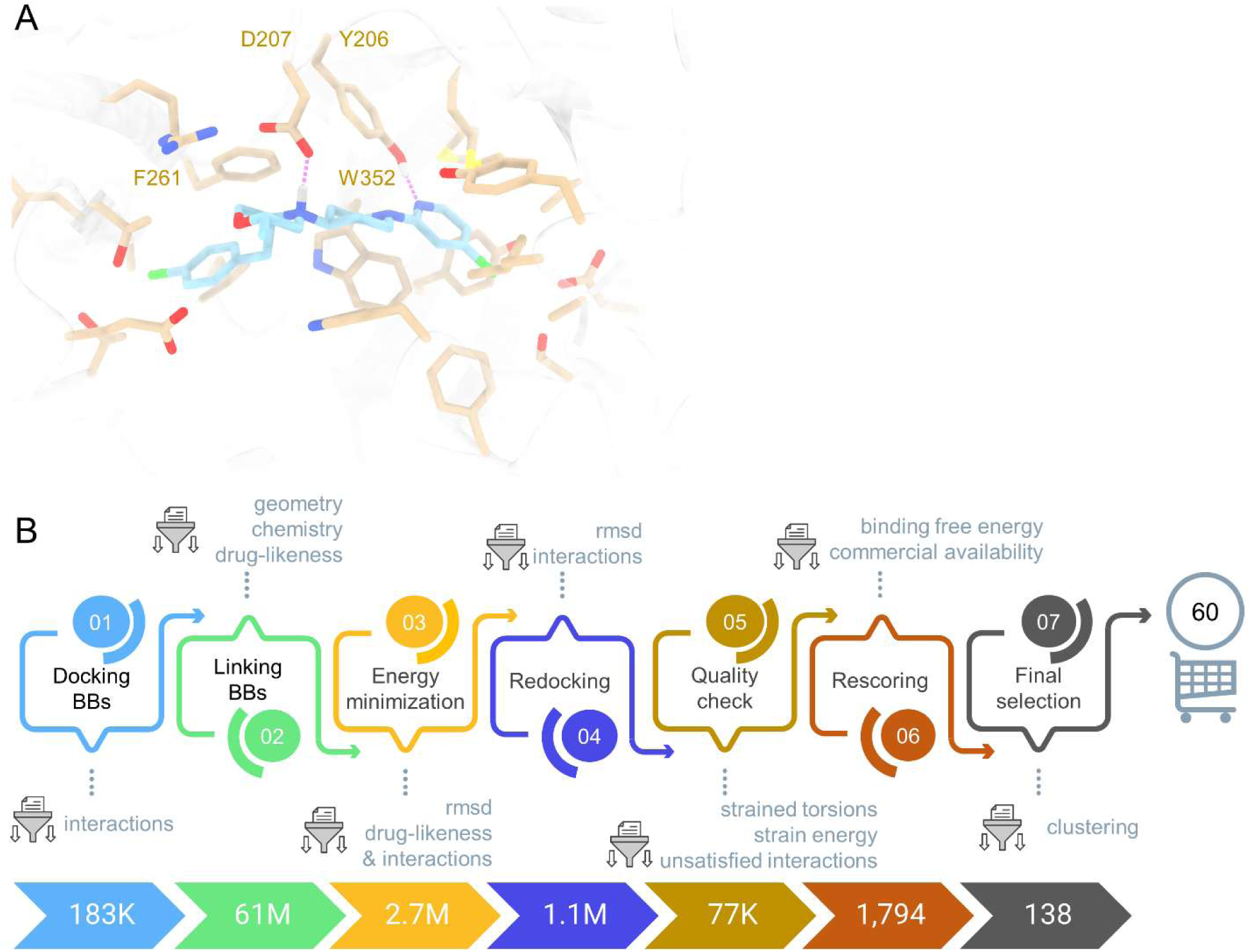
Structure-based organic-chemistry driven ligand design. **A**) Three-dimensional structure of CHI3L1 bound to a small molecule inhibitor (PDB 8R4X).^68^ Key protein-ligand hydrogen bonds are displayed by magenta dots. **B**) Generative design workflow yielding 60 synthesizable hits. At each step, several filters are introduced to restrain the generative design. We here ensured that the two key hydrogen bonds to Y216 and D217 are conserved throughout the workflow. Number of solutions are indicated, at each step, in the bottom diagram.

Reagent docking poses (20 per building block) are then systematically scanned for one of 40 possible organic chemistry linkages (**Table S1**), depending on strict topological (distances, angles, torsional angles) relationships and organic chemistry cross-compatibility. Upon successful linkage (covalent bond formation, ring closure), newly enumerated SpaceDock ligands are energy minimized in the target active site and redocked to ascertain that the same pose can be recovered again. Quality check of remaining poses^74^ eliminated dubious solutions (high strain energy, to many unsatisfied hydrogen-bonds and ionic bonds at the protein-ligand interface) which are then submitted to two independent and rigorous physics-based absolute binding free energy estimates (Hyde^75^ and VM2^76^) to keep only best scorers. Clustering by maximum common substructures afforded 60 clusters out of which the best VM2 scorer was purchased for synthesis at Enamine (Kyiv, Ukraine). Out of the 60 virtual hits, 45 could indeed be synthesized at sufficient purity (> 90% by LC-MS) to be tested experimentally.

### Primary screening by microscale thermophoresis

Virtual hits were tested at the single concentration of 250 µM for binding to a fluorescent-labelled His-tagged hCHI3L1 protein. Based on the normalized fluorescence (Fnorm) values (**Figure 2A**), 15 potential hits (**1c**, **1d**, **1e**, **2a**, **2d**, **2e**, **3a**, **3b**, **4a**, **4c**, **6a**, **7d**, **8a**, **8b**, **9e**) were identified. Comparison of compound fluorescence with a blank/reference (CHI3L1 labeled and treated with 2.5% DMSO instead of a test compound) identified eight compounds (**2d**, **2e**, **3a**, **3b**, **4c**, **6a**, **8b**, and **9e**) that exhibited higher or lower fluorescence than the blank (**Figure 2B**). To eliminate false positives due to signal interference or fluorophore interactions, additional control experiments were performed on those eight hits, including autofluorescence screening (**Figure 2C**) and a quenching assay (**Figure 2D**).

**Figure 2.**
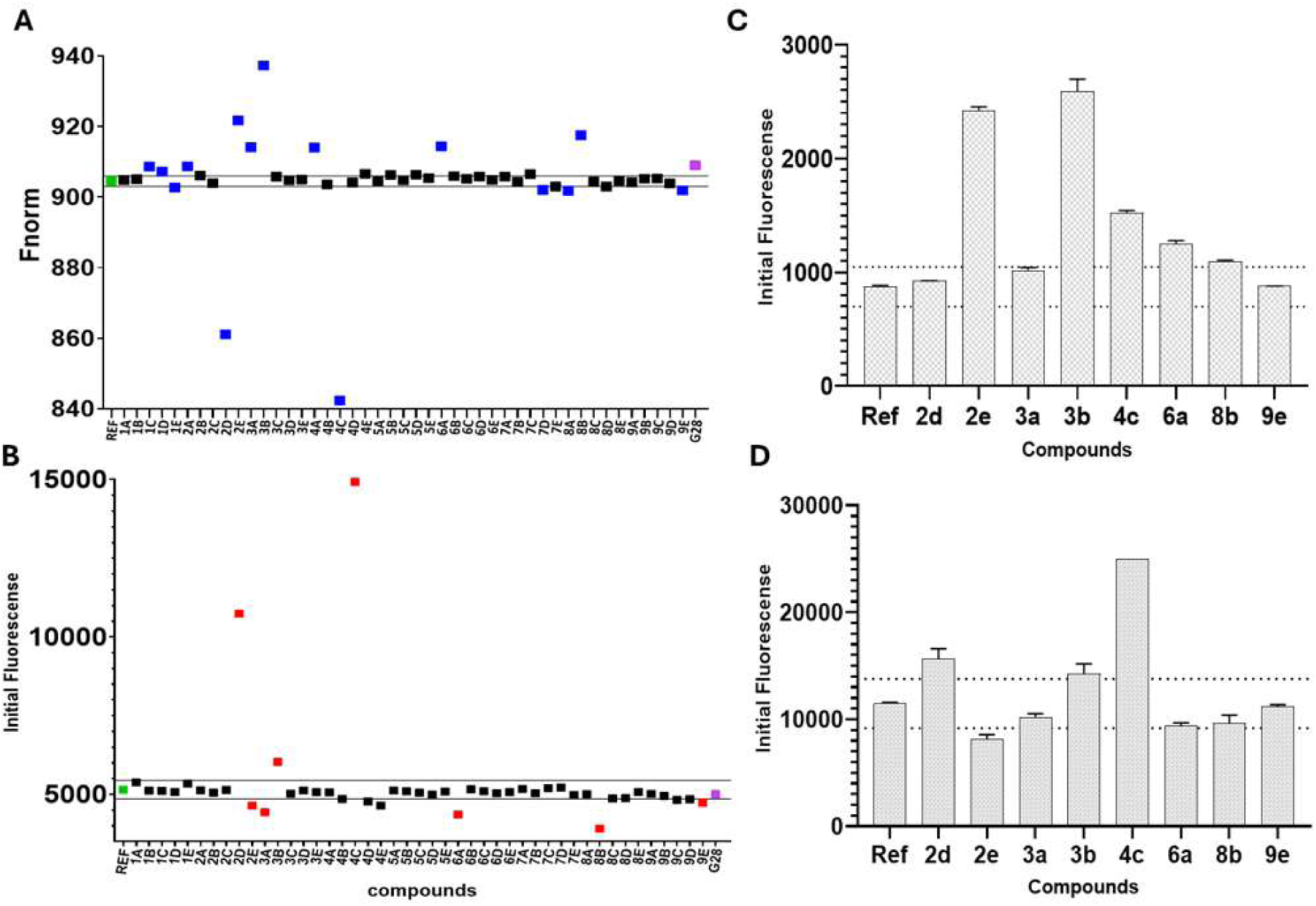
**(A)** Primary screening of molecules from at 250 μM (with 2.5% DMSO). The blue, green, black, and purple indicates the potential hits, negative control/reference (buffer with 2.5% DMSO), non-binder, positive control (**G28**)^67^; **(B)** Comparison of initial fluorescence of compounds with reference having labeled hCHI3L1 with 2.5% DMSO. The red dots indicate the compounds that may have signal interference or fluorophore interactions; **(C)** Autofluorescence and **(D)** quench test of candidates from single-dosage screening. Dotted lines in Figures 2C and 2D indicate the threshold values (mean negative control ± 3×SD) used to flag compounds with excessive autofluorescence or quenching.

In the autofluorescence assay, compounds **2e**, **3b**, **4c**, **6a** and **8b** exhibited high intrinsic fluorescence at the detection wavelength, whereas the fluorescence signals for compounds **2d**, **3a**, and **9e** remained within acceptable limits for MST analysis. In the quenching assay, compounds **2d**, **2e**, **3b**, and **4c** showed concentration-dependent quenching effects and were therefore excluded from further analysis.

Based on these criteria, compounds showing minimal signal interference across all assays were considered suitable for MST analysis. Through this refinement process, nine compounds—**1c**, **1d**, **1e**, **2a**, **3a**, **4a**, **7d**, **8a**, and **9e**—were confirmed as valid hits with reliable binding profiles.

### Dose-response confirmation

The potential hits identified from the single-dose screening were further evaluated in a dose-dependent MST assay to confirm their direct binding affinity for CHI3L1 (**Figure 3**). Among the nine candidates, three compounds **3a**, **4a**, and **9e** exhibited a clear dose-dependent response, indicating these are likely true binders.

**Figure 3.**
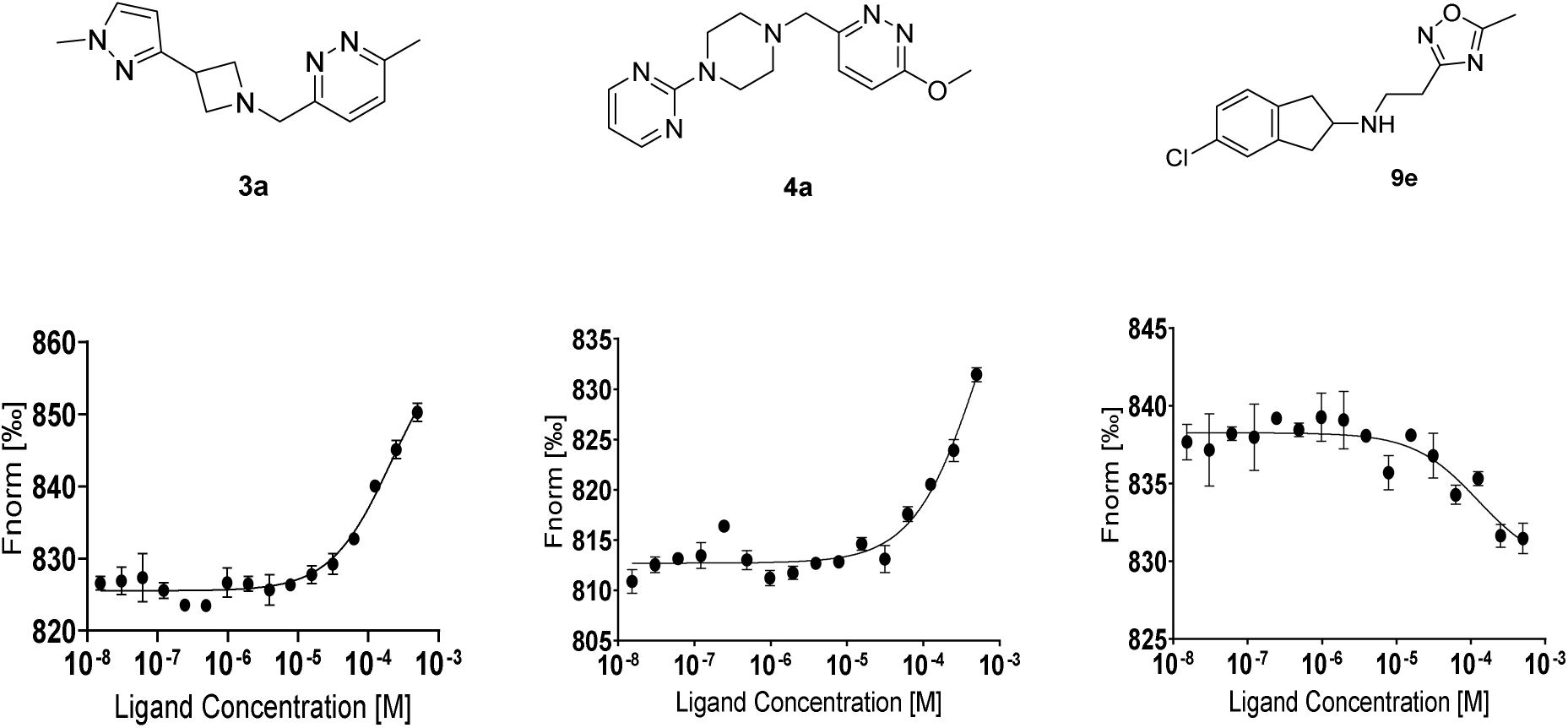
MST-based confirmation of CHI3L1 binding by compound **3a**, **4a** and **9e**, respectively. The graph displays dose-dependent changes in normalized fluorescence (Fnorm [%]) plotted against increasing concentrations of each ligand.

Interestingly, none of the three hits were available in the catalogue of 20 million commercially available screening compounds directly purchasable at E-molecule (https://www.emolecules.com/products/screening-compounds), thereby confirming the value of screening much larger “on-demand” chemical spaces.

### Hit confirmation by Surface Plasmon Resonance

Plasmon Resonance (SPR) was next employed to validate the binding interactions of selected compounds with CHI3L1 under label-free conditions. As a highly sensitive technique that enables real-time monitoring of molecular interactions, SPR serves as a valuable complement to microscale thermophoresis (MST). To support the MST results, we evaluated the binding affinities of compounds **3a**, **4a**, and **9e** using SPR. Compound **3a** (**Figure 4A**) did not exhibit a dose-dependent response, while compound **4a** (**Figure 4B**) showed a high relative response indicative of nonspecific binding. However, compound **9e** showed a clear, dose-dependent binding response, yielding a dissociation constant (K_d_) of 19.11 μM (**Figure 4C**).

**Figure 4.**
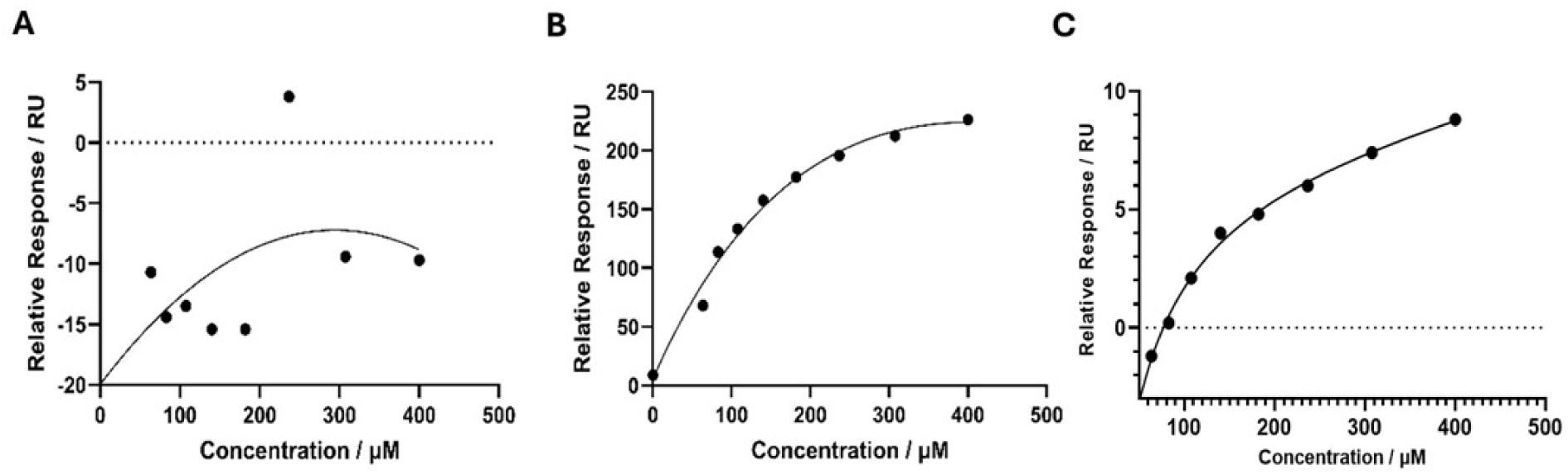
SPR-based binding affinity analysis of compounds **3a** (A), **4a** (B) and **9e** (C) towards hCHI3L1.

Given the low molecular weight of compound **9e** (MW=277.75) and an estimated logP of 2.31 (ACD/ChemSketch v2010), it presents an excellent ligand efficiency^84^ of 0.35 kcal mol^-1^ and a valuable start for hit to lead optimization. According to SpaceDock, the compound is hydrogen-bonded to two central polar residues (D207, Y206) and exhibit aromatic interactions to F261 and Y352. Distal aromatic groups (chloroindane, oxadiazole) fit into the two apolar ends of the pocket (**Figure 5**). However, the pocket remains empty at the vicinity of E290 and M204/M350, suggesting that adequate substitutions to reach both amino acids may improve the hit potency.

**Figure 5:**
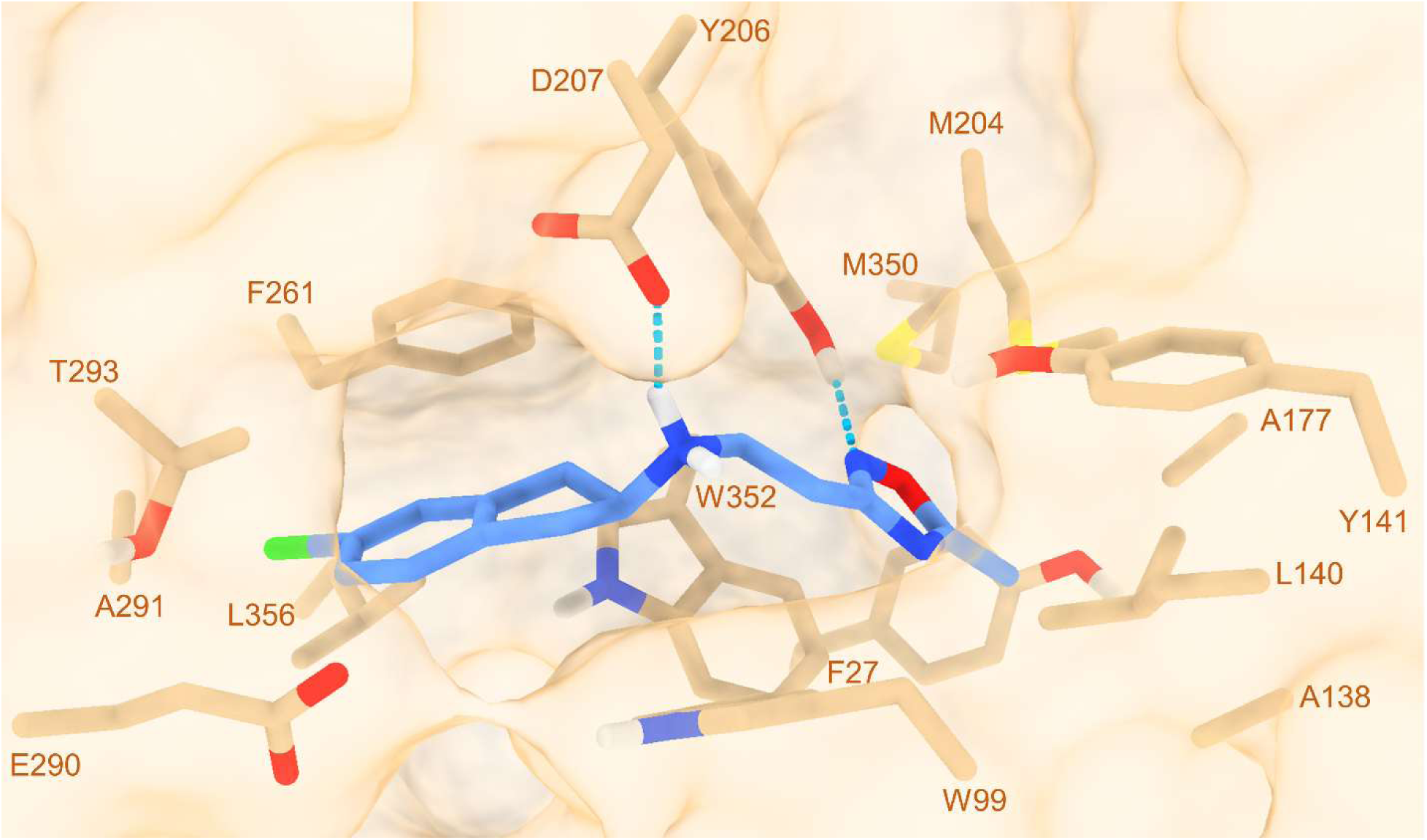
Proposed binding mode of compound **9e** (blue sticks) to human CHI3L1 (tan surface). Protein residues lining the ligand-binding pocket are labelled at the Calpha atom.

## AlphaLISA-Based Assessment of Compounds Targeting hCHI3LA–Galectin-3 Interaction

To investigate the ability of selected compounds to inhibit the interaction between hCHI3L1 and Galectin-3, compounds **3a**, **4a**, and **9e** were evaluated using an AlphaLISA-based inhibition assay (**Figure 6A**). The assay was performed in technical triplicates under optimized conditions. Compound **3a** exhibited no detectable inhibition, while compound **4a** showed a modest inhibition of 13.23%. Compound **9e** displayed the highest level of inhibition among the three, at 20.87%. These results suggest that **4a** and **9e** may weakly interfere with the hCHI3L1–Galectin-3 interaction, highlighting them as potential starting points for further optimization.

**Figure 6.**
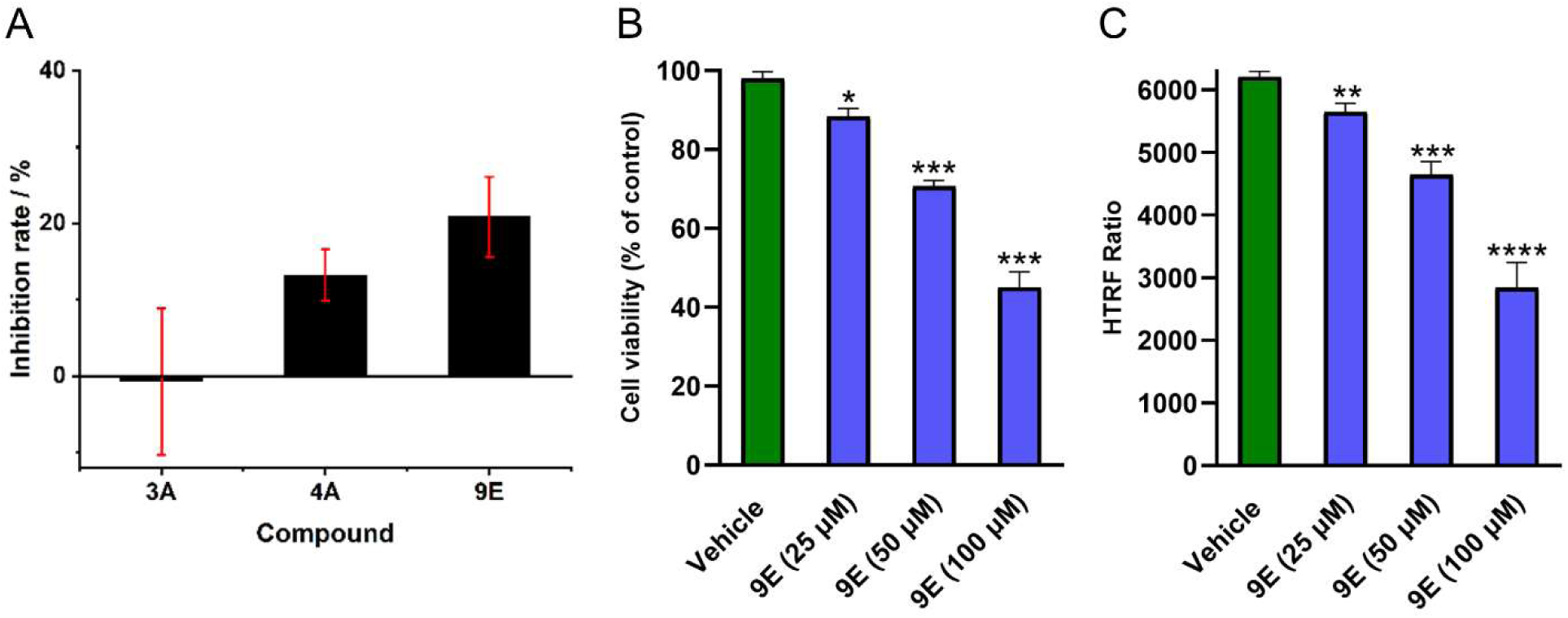
Experimental validation of compound 9e by orthogonal assays. (A) AlphaLISA-Based Inhibition of hCHI3L1-Galectin-3 Interaction by Compounds **3a**, **4a**, and **9e** at a 100 µM concentration; (B) Cell viability of GBM spheroids (as % of untreated control) upon incubation with increasing concentrations of compound **9e** after 72 h incubation. * *p* < 0.05 and *** *p* < 0.001, relative to vehicle control. Data are representative of three independent experiments. (C) Reduction in phospho-STAT3 levels, as determined by an HTRF phospho-STAT3 assay in GBM spheroids upon 72 incubation with compound **9e**. ** *p* < 0.01 and *** *p* < 0.001 relative to vehicle control. Data are representative of three independent experiments.

### Evaluation of the therapeutic potential using GBM organoids

To evaluate the functional impact of compound **9e** in a biologically relevant context, we utilized a 3D multicellular GBM spheroid model comprising GBM cells, endothelial cells, and macrophages. This model was chosen due to its ability to better recapitulate the complex tumor microenvironment characteristic of GBM. Treatment with compound **9e** led to a pronounced, dose-dependent decrease in spheroid viability, achieving statistical significance at concentrations as low as 25 μM (**Figure 6B**). This outcome aligns with the respective CHI3L1 binding affinity of compound **9e**. To further probe target engagement, we quantified phosphorylated STAT3 (pSTAT3) levels in GBM spheroids using a homogeneous time-resolved fluorescence (HTRF) assay. Compound **9e** elicited a dose-dependent reduction in pSTAT3 (**Figure 6C).** Collectively, these findings support compound **9e** as an inhibitor of the CHI3L1–STAT3 axis, capable of impairing tumor spheroid viability and dampening downstream signaling in a relevant 3D co-culture system.

## Conclusion

This study demonstrates the effective application of a structure-guided, reaction-aware virtual screening strategy to identify small-molecule inhibitors of CHI3L1, a non-enzymatic and conformationally dynamic protein implicated in GBM progression. By navigating a virtual chemical space of 377 billion compounds using the SpaceDock platform, we prioritized and synthesized top candidates, ultimately identifying compound **9e** as a validated hit. Compound **9e** exhibited micromolar binding affinity, measurable inhibition of CHI3L1–Galectin-3 interaction, and robust activity in a physiologically relevant multicellular 3D GBM organoid model, where it reduced tumor spheroid viability and suppressed downstream STAT3 signaling. These findings not only position **9e** as a promising lead for further optimization but also establish SpaceDock as a versatile platform for targeting structurally elusive proteins. The integration of computational ligand design, biophysical screening, and functional validation in tumor-relevant models offers a powerful framework for accelerating drug discovery in GBM and other cancers where conventional approaches have proven insufficient.

## ACKNOWLEDGMENTS

The Calculation Centre of the IN2P3 (CNRS, Villeurbanne, France) is acknowledged for the allocation of computing time and excellent support.

ASSOCIATED CONTENT

## Supporting Information

The following files are available free of charge.

Organic chemistry reactions used to generate targets-specific ligands, SMILES strings of 45 virtual hits selected for synthesis and experimental validation, and LC-MS data for the top hit compounds (PDF).

AUTHOR INFORMATION

## Author Contributions

The manuscript was written through contributions of all authors. All authors have given approval to the final version of the manuscript.

## Notes

D.R. is cofounder and shareholder of BIODOL Therapeutics. The other authors declare no competing interests.

## Abbreviations

Gal3: Galectin-3
CHI3L1: Chitinase-3-like protein 1
HTRF: homogeneous time-resolved fluorescence homogeneous time-resolved fluorescence
IL-13Rα2: IL-13 receptor alpha 2
PPI: protein-protein interactions
MST: microscale thermophoresis
SPR: surface plasmon resonance
KD: equilibrium dissociation constant
RAGE: receptor for advanced glycation end products
VEGFR2: Vascular Endothelial Growth Factor Receptor.

---

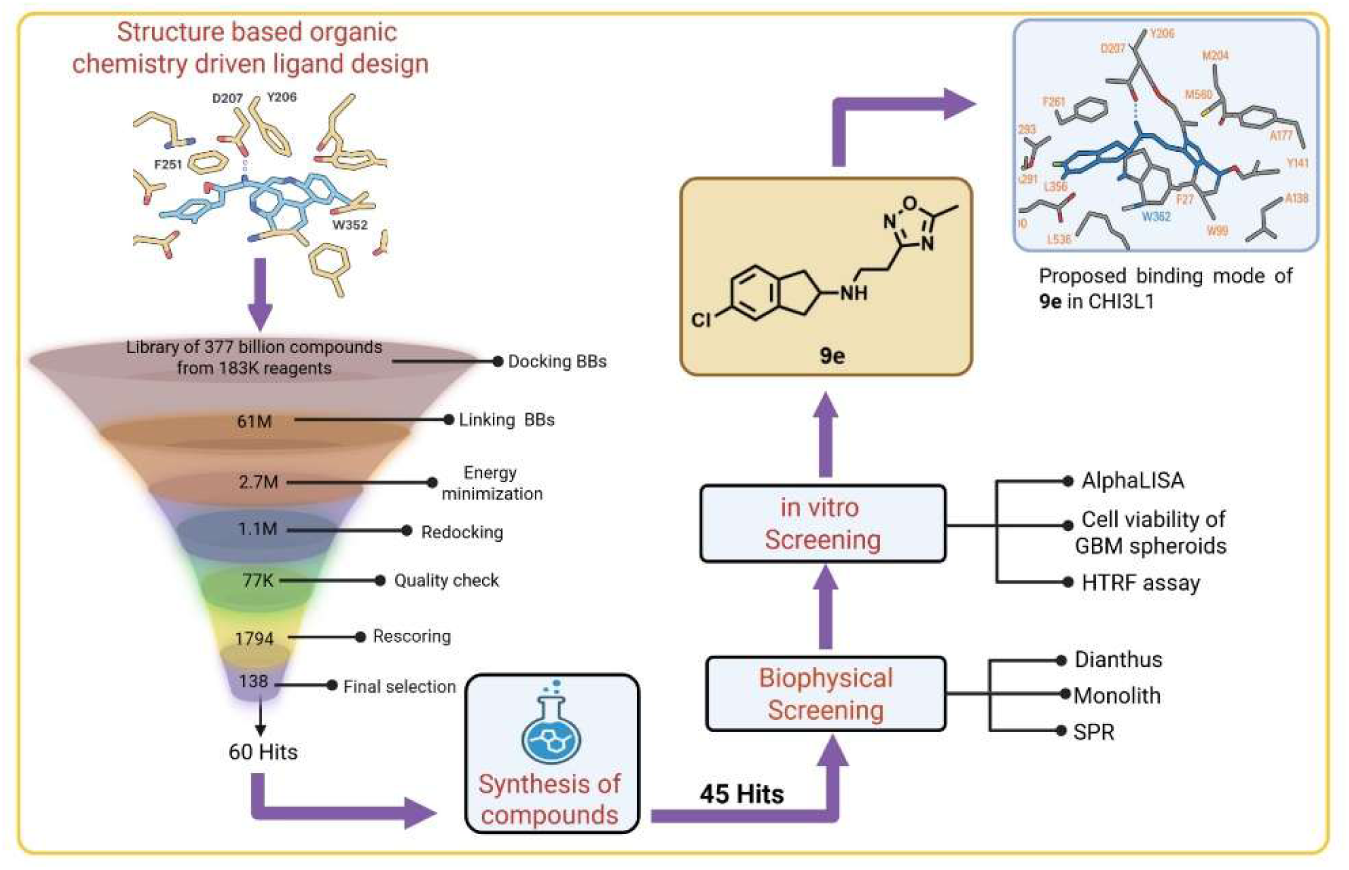
Table of Contents artwork

