## Supporting Information for "Structure-Guided Discovery of CHI3L1 Inhibitors from Ultralarge Chemical Spaces for Glioblastoma Therapy"

**Contents**

**Table S1.** Organic chemistry reactions used to generate targets-specific ligands	S2

**Table S2.** SMILES strings of 45 virtual hits selected for synthesis and experimental validation S3

**Figure S1.** LC-MS Characterization of Compound **3a**. S5

**Figure S2.** LC-MS Characterization of Compound **4a**. S5

**Figure S3.** LC-MS Characterization of Compound **9e**. S6

**Table S1. Organic chemistry reaction used to guide the generative ligand design**

| **Reaction** | **Products** |
| --- | --- |
| Amide bond | 2,080,278,830 |
| Benzimidazole | 12,133,689 |
| Benzofuran | 458,781 |
| Benzothiazole | 41,104 |
| Benzothiophene | 27,577 |
| Benzoxazole (from carboxylic acids) | 596,144 |
| Benzoxazoles (from aldehydes) | 6,034,080 |
| Buchwald-Hartwig | 702,785,160 |
| Decarboxylative coupling | 2145,240 |
| Double reductive amination | 370,619,894,598 |
| Fisher_indole | 421,139 |
| Friedlaender_quinoline | 36,257 |
| Grignard | 362,592,048 |
| Heck | 18,703,776 |
| Huisgen_Cu | 63,730,611 |
| Huisgen_Ru | 63,730,611 |
| Indole | 507,808 |
| Mitsunobu | 85,589,044 |
| MonoHetarylation | 247,984,956 |
| N_alkylation | 534,643,824 |
| N_arylation | 4,441,268 |
| Negishi | 753,337,809 |
| Niementowski_quinazoline | 2,686,761 |
| Oxadiazole | 144,510,240 |
| Phthalazinone | 27,630 |
| Pictet_Spengler | 1,848,681 |
| Reductive amination | 446,196,836 |
| SNAr | 235,734,140 |
| Sonogashira | 28,955,304 |
| Spirochromanone | 17,889 |
| Stille | 1,949,040 |
| Sulfonamide | 95,586,432 |
| Sulfone | 546,390,625 |
| Sulfonic_esters | 3,969,343 |
| Suzuki | 35,213,308 |
| Thiazole | 188,594 |
| Thiourea | 16,241,890 |
| Triazole | 154,186,216 |
| Urea | 42,391,960 |
| Williamson-Ether | 92,592,591 |
| **TOTAL** | **377,408,814,932** |

**Table S2.** SMILES strings of 45 virtual hits selected for synthesis and experimental validation

| **Manuscript ID** | **Catalog ID** | **Smiles** |
| --- | --- | --- |
| 1A | Z7352864808 | FC=1C=CC=C(C1)OC2CCN(CC3=NOC=C3Cl)C2 |
| 1B | Z3390876489 | CC=1N=COC1CN2CC(C2)N3C=C(Br)C=N3 |
| 1C | Z9429368575 | IC=1C=NC(CN2CCC=3NC(=NC3C2)C=4C=CC=CC4)=NC1 |
| 1D | Z9429368666 | CC=1C=CC(=CN1)C2(O)CCN(CCOC=3C=CC=CC3)C2 |
| 1E | Z3877275693 | CC=1C=NN(C1)C2CCN(CC3=NOC(C)=C3Br)C2 |
| 2A | Z4031787257 | C=1C=CC=2NC(CN3CCCC3CC=4C=CC=NC4)=NC2C1 |
| 2B | Z4275701597 | OC1(CCN(CCN2C=C(Cl)C=N2)C1)C=3C=CC=NC3 |
| 2C | Z5593737578 | ClC=1C=CC(=NC1)C2CN(CCN3C=NC(Br)=N3)CCO2 |
| 2D | Z3390875919 | OC=1C=CC=NC1CN2CC(C2)N3C=C(Br)C=N3 |
| 2E | Z5872853768 | CC=1C=NC=C(C1)C2CCN(CC=3N=CC=CC3O)C2 |
| 3A | Z4429787323 | CC=1C=CC(CN2CC(C2)C=3C=CN(C)N3)=NN1 |
| 3B | Z3520346581 | CC=1C=C(ON1)C2CCN(CC3=CC=NO3)C2 |
| 3C | Z9429368611 | CC1=NOC(=N1)C2CCN(CCC(=O)NC=3C=CC=CN3)C2 |
| 3D | Z3321740114 | CC1=NC(=CS1)CCN2CCC(O)(C2)C=3C=CC(C)=CC3 |
| 3E | Z9429368653 | CC1=CON=C1CN2CC(C2)C=3N=CC(Br)=CN3 |
| 4A | Z2015085911 | COC=1C=CC(CN2CCN(CC2)C=3N=CC=CN3)=NN1 |
| 4B | Z5069420319 | CC1=CNC(CN2CC(C2)C3=NN(C)C=C3F)=N1 |
| 4C | Z7564998071 | CCOC=1C=CC(=NC1)C(C)NCC=2N=CC(I)=CN2 |
| 4D | Z9429368752 | C#CC=1C=NC=C(C1)CNC[C@H](O)C=2C=CC(C)=CC2 |
| 4E | Z2179944226 | N#CC=1C=CC(=NN1)NCCC2=NC=C(Br)S2 |
| 5A | Z2473812849 | OC1(CCCN(CCN2C=C(Br)C=N2)C1)C=3C=CC(Cl)=CC3 |
| 5B | Z9429368696 | CC1=CC(=NO1)NC(=O)CCN2CC(C)C(C2)C=3C=CC(Br)=CC3 |
| 5C | Z1653302338 | CN1C=NN=C1CN2CCC(C2)OC=3C=CC=C(F)C3 |
| 5D | Z2202848624 | CC=1C=C(CNC2CCC(CC2)C3=NC(C)=NO3)ON1 |
| 5E | Z4207100469 | BrC=1C=C(CNC2CCN(CC2)C=3N=CC=CN3)ON1 |
| 6A | Z5398100628 | CC=1C=CC(CN2CC(C2)N3N=CC(Br)=N3)=NN1 |
| 6B | Z5873022783 | CC1CN(CC2=NN=C(O2)N(C)C)CC1C=3C=CC(Br)=CC3 |
| 6C | Z5646543175 | C#CC=1C=CC=2CCN(CCN3C=C(C)C=N3)CC2C1 |
| 6D | Z3315951091 | CN1C=C(N=N1)C2CCCN(CCC=3C=NC=C(F)C3)C2 |
| 6E | Z5646695083 | COC=1C=C(CN2CCC(CC2)C=3N=NN(C)N3)ON1 |
| 7A | Z9429368649 | CC1=NC(=NN1)C2CN(CC3=CN=C(S3)N(C)C)C2 |
| 7B | Z9429368590 | CN1C=NN=C1CN2CC(C2)C=3N=CC(Br)=CN3 |
| 7C | Z5032469301 | CC1=CC=C(O1)C2CN(CCC=3C=C(C)ON3)CCO2 |
| 7D | Z9429368724 | CC(C=1C=CC=2OCCCOC2C1)N(CC=3N=CC(I)=CN3)C4CC4 |
| 7E | Z1130197780 | FC=1C=CC(=CC1)OCCN2CCC=3NC=4C=CC=CC4C3C2 |
| 8A | Z7488224760 | CC1=CN=C(CN2CCC=3N=C(Br)SC3C2)O1 |
| 8B | Z5638578884 | CC1=NN=C(CN2CCC(CC2)C=3N=NN(C)N3)S1 |
| 8C | Z9429368595 | CN1C=C2CCC(CC2=N1)NCC3=CC=4SC=C(Br)C4N3 |
| 8D | Z5090083629 | CC=1C=NN(C1)CCN2CCOC(C2)C=3C=CC(C)=CC3 |
| 8E | Z7513014071 | CN1N=NC(=N1)C2CCCN(CCN3C=C(Br)C=N3)C2 |
| 9A | Z4155591198 | CCC=1ON=C(CN2CCC(C2)C3=NC(C)=NO3)C1C |
| 9B | Z3390869876 | CC1=NC(=NO1)C2CN(CC=3C=C(NN3)C=4C=CC=CC4)C2 |
| 9C | Z4271867511 | COCC1=NN=C(CN2CCOC3C=4C=CC=CC4OCC32)S1 |
| 9D | Z7058219934 | CC1=NC(=NO1)C2CCN(CCOC=3C=NC=C(Cl)C3)C2 |
| 9E | Z1607704570 | CC1=NC(CCNC2CC=3C=CC(Cl)=CC3C2)=NO1 |

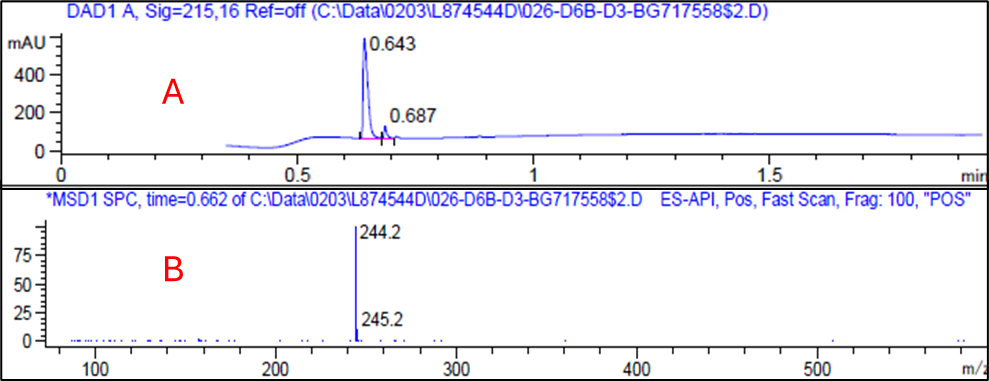

**Figure S1.** LC-MS Characterization of Compound **3a**. **(A)** Total Ion Chromatogram (TIC) showing a sharp peak at 0.643 min, indicating the retention time of Compound **3a**; **(B)** Mass spectrum in positive ionization mode (ESI+), calcd. [M+H]^+^ 244.3.

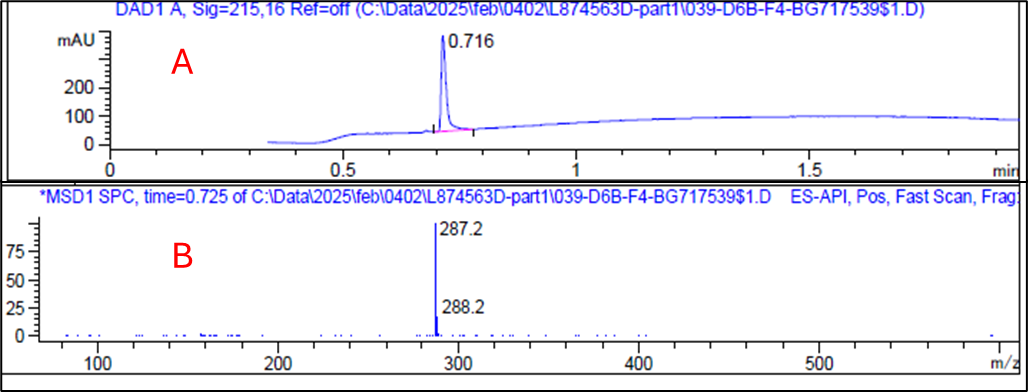

**Figure S2.** LC-MS Characterization of Compound **4a**. **(A)** Total Ion Chromatogram (TIC) showing a sharp peak at 0.716 min, indicating the retention time of Compound **4a**; **(B)** Mass spectrum in positive ionization mode (ESI+), calcd. [M+H]^+^ 287.3.

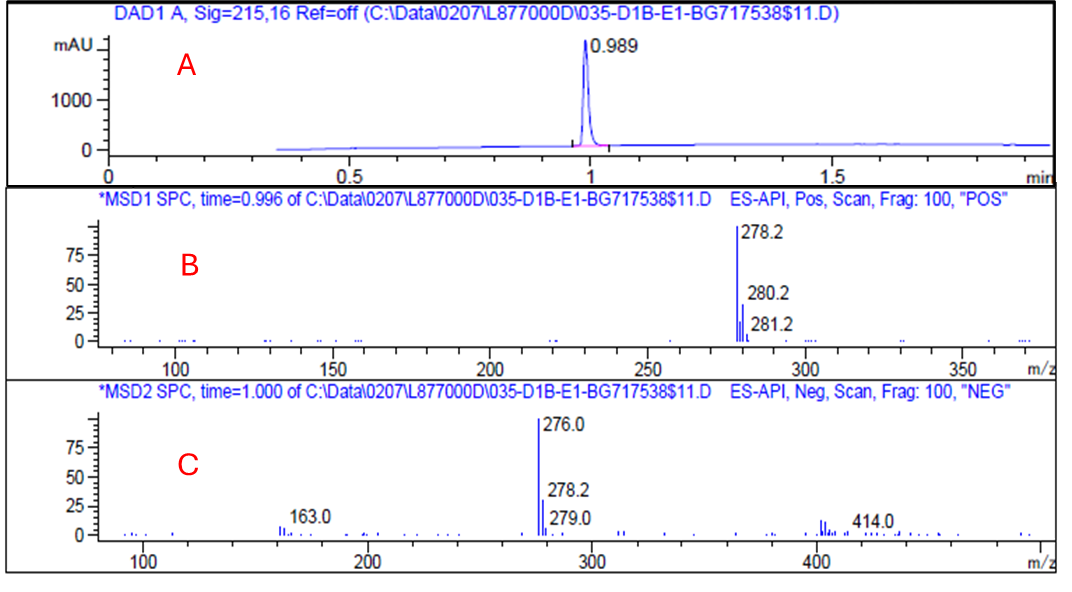

**Figure S3.** LC-MS Characterization of Compound **9e**. **(A)** Total Ion Chromatogram (TIC) showing a sharp peak at 0.999 min, indicating the retention time of Compound **9e**; **(B)** Mass spectrum in positive ionization mode (ESI+), calcd. [M+H]^+^ 278.1; **(C)** Mass spectrum in negative ionization mode (ESI–).
